# Serum biochemistry suggests grey squirrels (*Sciurus carolinensis*) have poorer physiological condition in urban settings

**DOI:** 10.1101/2019.12.16.878702

**Authors:** Chloé Schmidt, Jason R Treberg, Riikka P Kinnunen, Colin J Garroway

## Abstract

Human food waste in cities presents urban wildlife with predictable, easily accessible high-calorie food sources, but this can be both beneficial and harmful for individual health. We analyzed body condition and serum chemistry (electrolyte levels, markers of kidney and liver function, protein, glucose, and cholesterol) in an urban and rural population of eastern grey squirrels (*Sciurus carolinensis*) to assess whether proximity to the human food waste that is associated with urban habitats had ill effects on health. We found no differences in body condition between habitats and no evidence of malnutrition at either site. However, urban squirrels had higher blood glucose, lower potassium, phosphorus, chloride, and albumin:globulin ratios. These results align with previous findings of increased dietary sugar in cities, and suggest that urban populations of grey squirrels are under greater environmental stress than rural populations.

## Introduction

The distribution of resources available to animals is fundamentally altered in cities. Urban vegetation and tree cover, important natural food sources for wildlife, are reduced and patchy. Exotic species in urban plant communities may differ in phenology from native species, leading to changes in diet and foraging behavior among urban fauna. Human food waste is a plentiful new resource in cities which often allows animals to reach high population densities. Dietary shifts and higher densities in stressful urban environments could cause animals to be in poor physiological condition (1, 2). To explore how changing habitat quality associated with urbanization affects animal physiology, we looked for signs of ill health in a successful urban mammal, the eastern grey squirrel (*Sciurus carolinensis*).

In general, urban wildlife populations will lose access to natural food sources while gaining access to new foods associated with human trash and supplemental feeding. Human foods found as litter tend to be easily accessible, calorie-rich (3), and have lower nutritional value than natural food sources (4). Human food subsidies are a double-edged sword: increased calorie intake may positively influence body condition, survival, and reproductive success, however at the same time its limited nutrient quality may make it detrimental to animals’ overall health (5). Alterations in food availability, predictability, and quality has the potential to induce major shifts in nutritional status among urban wildlife (3).

Successful urban colonizers tend to be generalist, opportunistic omnivores that can take advantage of human food waste, sometimes to the point of complete dependency (6). In such species, urbanization leads to rapid changes in diet composition (2, 3, 6). Diets high in carbohydrates and saturated fats are linked to increasing trends in obesity in humans (7), as well as to elevated risk of cardiovascular disease, Type 2 diabetes and other metabolic disorders (8, 9). Similar metabolic conditions could also occur in non-human wild species (10–12). For instance, urban crows have been shown to have elevated plasma cholesterol (13), and brown bears with access to human food subsidies have higher carbohydrate and lower protein levels than those in more natural habitats (14). Greater body mass in urban-dwelling individuals has been reported in rats (15), foxes (16, 17), baboons (18), deer (19), and raccoons (20). Whether these symptoms indicate poor health is uncertain. Urban racoons were found to be heavier and hyperglycemic, but leptin levels (which are higher in obese humans) were normal (20). Urban baboons with access to human garbage exhibited symptoms consistent with metabolic disorder in humans such as high body mass index, hyperglycemia, high leptin, and insulin resistance (18). For the most part however, we do not know if the risk of developing metabolic disorders should be a general concern among urban mammals (2).

We compared results from a suite of standard biochemical assays of blood samples collected from urban and rural eastern grey squirrels. Serum biochemistry analysis is an effective way to measure population-level health, as it is a good indicator of disease state, nutritional status, and habitat quality (21). If we assume that rural individuals living in natural environments generally have normal, approximately healthy blood biochemistries, then we can treat those results as a baseline from which deviations in urban populations could point to health issues. Grey squirrels are a successful urban species, abundant in cities both within and outside their natural range (22). They are omnivorous with a wide dietary breadth, typically consisting of the nuts, flowers, and buds of a variety of hardwood trees (such as oaks, hickory, and beech), fungi, insects, cultivated crops, and from time to time, small animals and bones (23). Grey squirrel abundance is positively associated with human food provisioning (24, 25). More anecdotally, grey squirrels are frequently observed feeding on high calorie human food waste (26–29) and on birdseed at feeders which is known to be less healthy than natural foods for birds (4). We measured body condition, and used serum chemistry analyses to assess electrolyte levels, markers of kidney and liver function, protein, glucose, and cholesterol, to get a general picture of population health in urban and rural squirrels. We hypothesized that, due to differing food sources, urban squirrels would show signs of ill health related to a poor-quality urban diet. Similar to previous studies of urban mammals, we specifically expected to find higher glucose, cholesterol, and body condition in urban squirrels.

## Methods

### Sampling

Grey squirrels were sampled from May – June 2019 in two sites with differing levels of anthropogenic food. The ‘urban’ site is an approximately 10 ha park located on the University of Manitoba campus in Winnipeg, Manitoba. Relative to natural forests it is sparsely treed, and is adjacent to a river, a suburb, and bordered by major roads on two sides. This park experiences high human foot traffic. Squirrels at this site have easy access to birdfeeders and human food waste in trash cans, and litter. Our second site was a rural hardwood forest patch of approximately 34 ha located in southern Manitoba (49°14’39"N, 98°00’47"W). The forest is bordered by agricultural land and a road (at a distance of 47m), and has minimal anthropogenic food sources.

Squirrels were trapped using Tomahawk live traps (Tomahawk Live Trap Co., Tomahawk, WI, USA) baited with peanut butter, then restrained in a capture bag (30) without anesthetic. Sex, approximate age (adult or juvenile), weight (grams), reproductive status (scrotal or non-scrotal for males; lactating or non-lactating for females), and body length (centimeters) from the base of the skull to the base of the tail were recorded for each individual. We implanted passive integrated transponder (PIT) tags for later identification. In total, 20 individuals were trapped at the urban location, and 10 in the rural location. We measured body condition by taking the residuals from a linear regression of mass on body size (here, spine length). This method corrects for the effects of an individual’s structural size on its mass (31, 32). We computed body condition independently for the sexes within each location. Juveniles were excluded from body condition calculations (Table 1). Squirrels were released after capture. Our sampling protocol was approved by the University of Manitoba animal care and use committee.

**Table 1.**
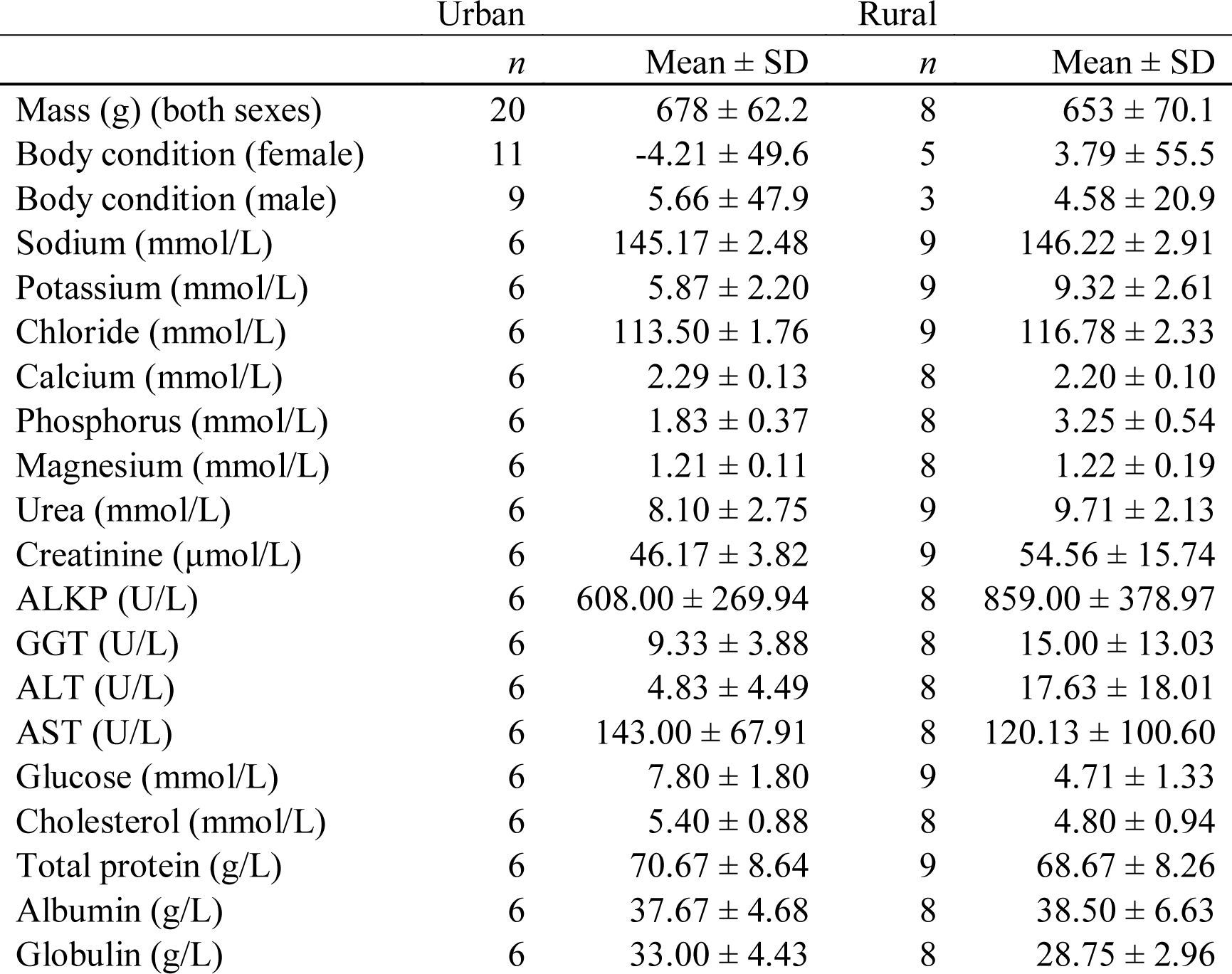
Body condition and serum parameters in urban and rural grey squirrels. Sample sizes differ when individuals were excluded due to age (mass and body condition) or low serum sample volume.

### Serum samples and tests

Blood samples were collected from a subset of all individuals captured at each site (Table 1). Samples were taken at the urban site between May 13-15, 2019, and the rural site between June 7-14, 2019. Blood (≤1 mL) was drawn on-site from the femoral vein using a syringe without anticoagulant and stored on ice until delivery to the Manitoba Veterinary Diagnostics Services lab (department of Manitoba Agriculture Food and Rural Development, Winnipeg, MB). Samples were processed within 12h of capture. Serum biochemistry profiles were run for each sample, consisting of the following tests: sodium, potassium, chloride, urea, creatinine, calcium, phosphorus, magnesium, amylase, lipase, alkaline phosphatase (ALKP), gamma-glutamyl transferase (GGT), bilirubin, alanine aminotransferase (ALT), aspartate aminotransferase (AST), creatine kinase, glucose, cholesterol, total protein, albumin, and globulin. Some tests were omitted when sample volume was lacking; for this reason, we do not report results for amylase, lipase, or creatine kinase. Sample sizes for each comparison are given in Table 1.

### Statistical analysis

Boxplots were created in R version 3.6.1 (33). We first plotted and visually inspected the data, then used principal components analysis (PCA) to visualize groupings, if any. We then tested differences in serum variables between sites using non-parametric Kruskal-Wallis tests due to small sample size and unbalanced groups. We found no sex differences for serum measurements (Fig. 1, 2, 3), thus did not include sex as a factor in our analyses.

**Figure 1.**
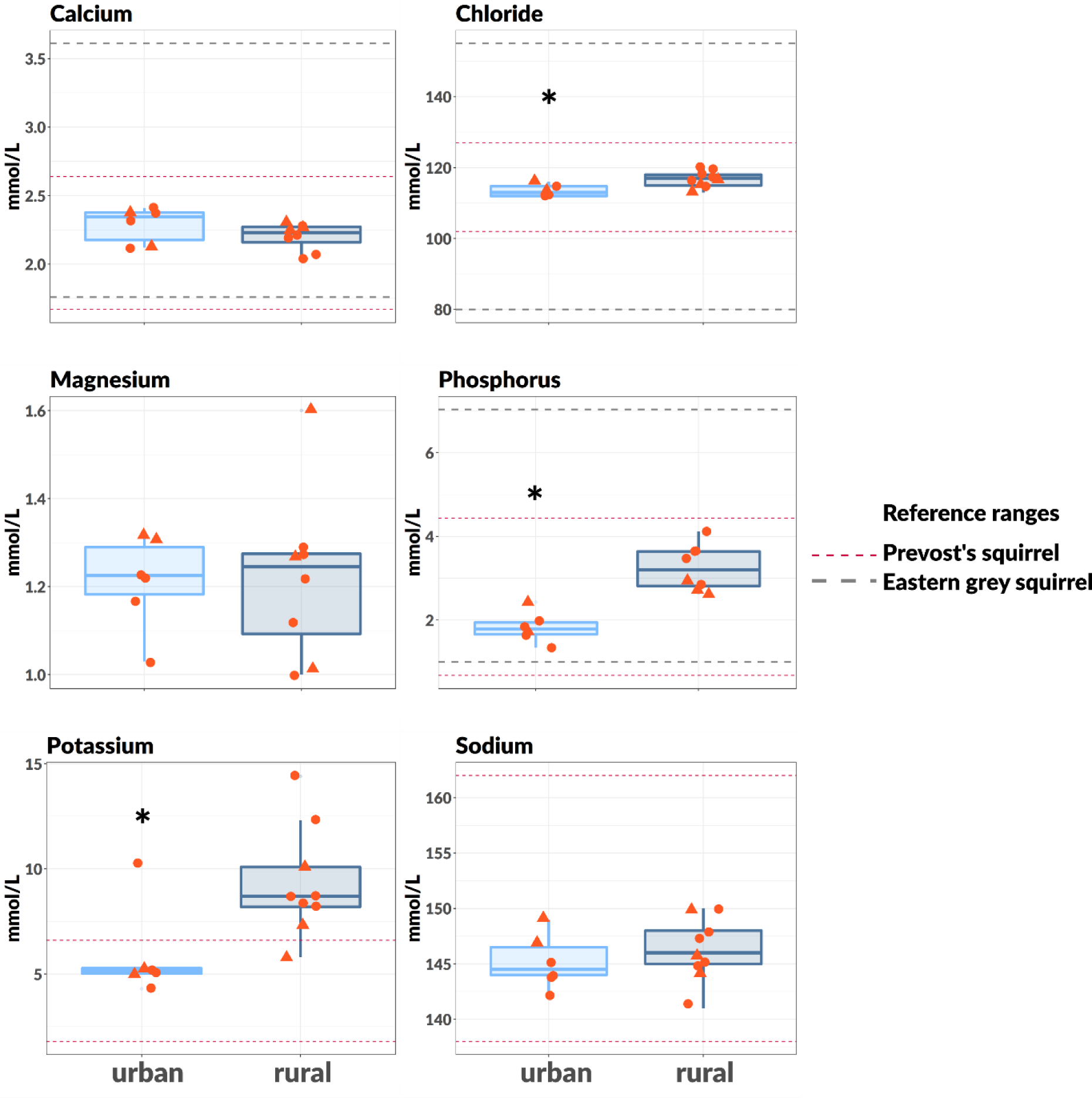
Boxplots for serum electrolytes. Circles are females, triangles are males. Reference ranges are shown for analytes when available. Asteriks (*) indicate noteworthy differences between urban and rural populations (*p* < 0.05).

**Figure 2.**
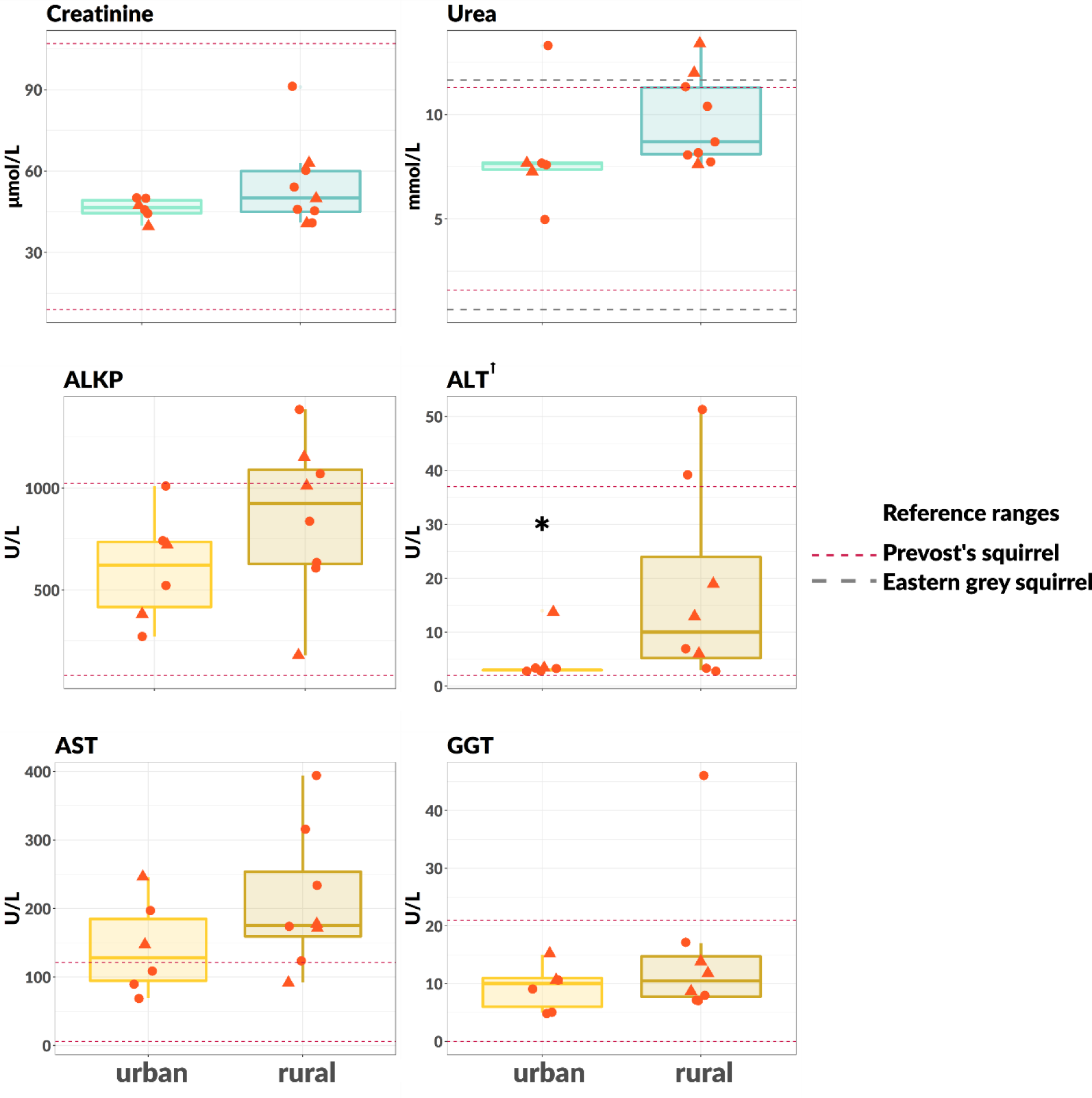
Boxplots for creatinine, urea, and enzymatic markers for liver and kidney function. Circles are females, triangles are males. Reference ranges are shown for analytes when available. ALKP = Alkaline phosphatase; ALT = alanine aminotransferase^Ɨ^; AST = aspartate aminotransferase; GGT = gamma-glutamyl transferase. Asteriks (*) indicate noteworthy differences between urban and rural populations (*p* < 0.05). ^Ɨ^Note: 5 of 6 urban samples had ALT levels below the detectable limit (3 U/L).

**Figure 3.**
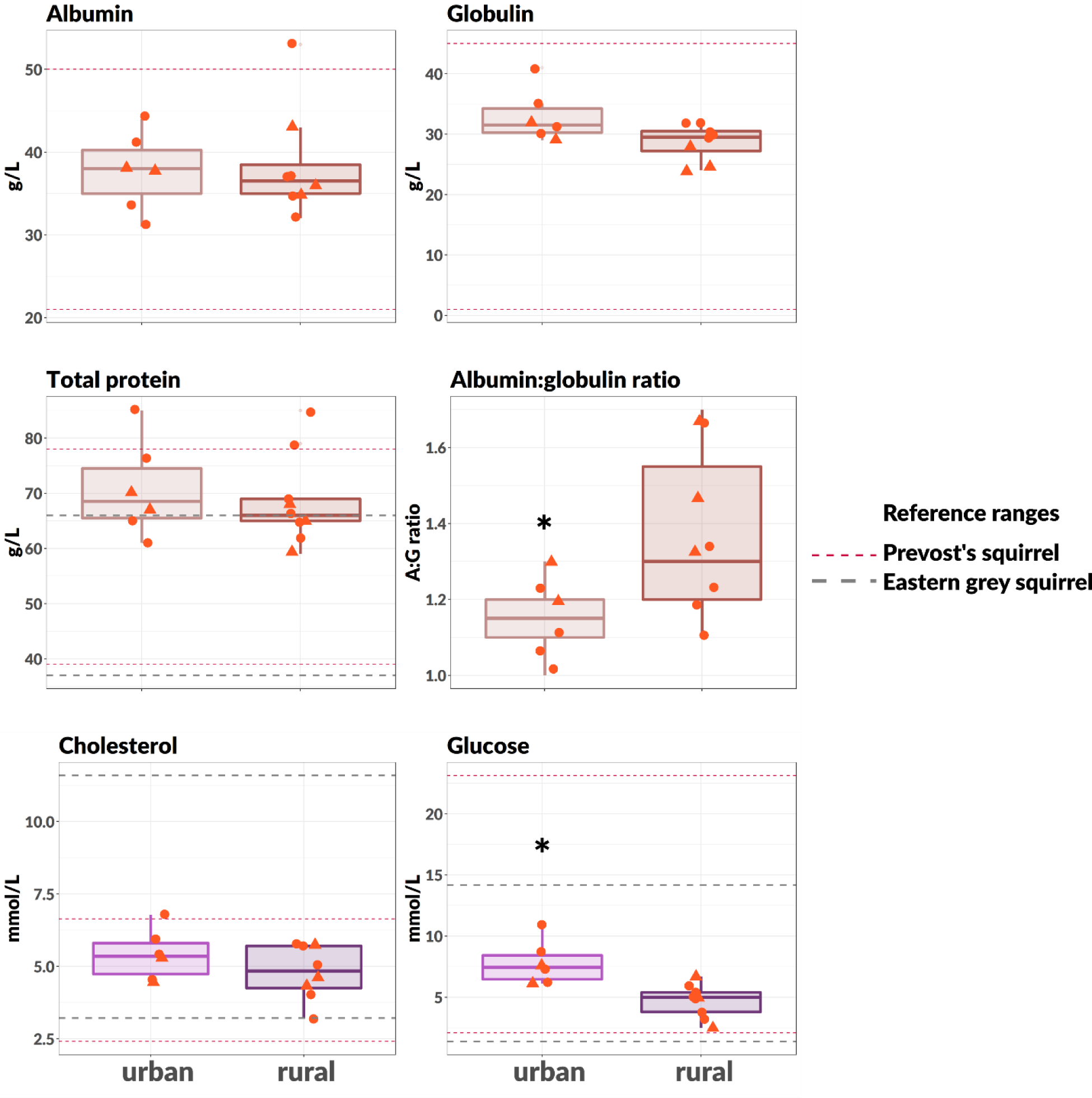
Boxplots for serum protein, cholesterol, and glucose. Circles are females, triangles are males. Reference ranges are shown for analytes when available. Asteriks (*) indicate noteworthy differences between urban and rural populations (*p* < 0.05).

## Results

Mean body mass (urban: 678 ± 62.2, rural: 653 ± 70.1, mean ± SD), and body condition (urban: 0.23 ± 47.8, rural: 4.09 ± 43.4) for all adult individuals trapped between May-August were similar between sites (Table 1). There were no detectable differences in body condition between sexes (Table 1).

Urban squirrels had higher glucose levels than those from the rural site (*P* <0.01, *X*^2^ = 8.70; Fig. 1, Table 1), in line with our expectations. Cholesterol levels did not detectably differ by location (Fig. 1, Table 1). Among serum ions, potassium (*P* = 0.02, *X*^2^ = 5.57), phosphorus (*P* < 0.01, *X*^2^ = 9.6), and chloride (*P* = 0.01, *X*^2^ = 5.95) were lower in urban squirrels (Fig. 1, Table 1). Sodium, calcium and magnesium levels were comparable at both sites (Fig. 1, Table 1).

ALT was lower at the urban site (*P* = 0.05, *X*^2^ = 3.73; Fig. 2, Table 1). A majority of urban samples were below the detection limit (3 U/L) (Fig. 2). We did not detect any differences in other serum enzymes (ALKP, GGT, AST; Figure 2). Total bilirubin, creatinine, urea, total protein levels, as well as albumin and globulin were similar at both sites (Figs. 2 and 3, Table 1). The ratio of albumin to globulin (A:G ratio) was higher in rural squirrels (*P* = 0.05, *X^2^* = 3.91; Fig. 3).

Seven principal components had eigenvalues greater than 1—these accounted for 91% of the variation in response variables. PC1 (25.6% of total variation) captured the variation between sites (Fig. 4, Table 2). Chloride, phosphorus, glucose and AST loaded most strongly on PC1 (Table 2).

**Table 2.** Factor loadings on the first two principal components (PCs). Urban and rural sites were separated along PC1.

|  | PC1 | PC2 |
| --- | --- | --- |
| Sodium | 0.156365 | -0.27516 |
| Potassium | 0.277074 | 0.29981 |
| Chloride | 0.334378 | -0.1606 |
| Urea | 0.164741 | 0.338645 |
| Creatinine | 0.096411 | 0.055769 |
| Calcium | -0.29924 | 0.046322 |
| Phosphorus | 0.395276 | 0.08438 |
| Magnesium | 0.034263 | 0.180375 |
| ALKP | 0.228345 | 0.092981 |
| GGT | 0.200064 | 0.219328 |
| ALT | 0.256803 | -0.23847 |
| AST | 0.344987 | 0.005621 |
| Glucose | -0.379 | 0.008177 |
| Cholesterol | -0.2204 | 0.170467 |
| Total protein | -0.0298 | 0.465441 |
| Albumin | 0.069876 | 0.428948 |
| Globulin | -0.15862 | 0.316649 |
| Body condition | -0.05741 | 0.073088 |

**Figure 4.**
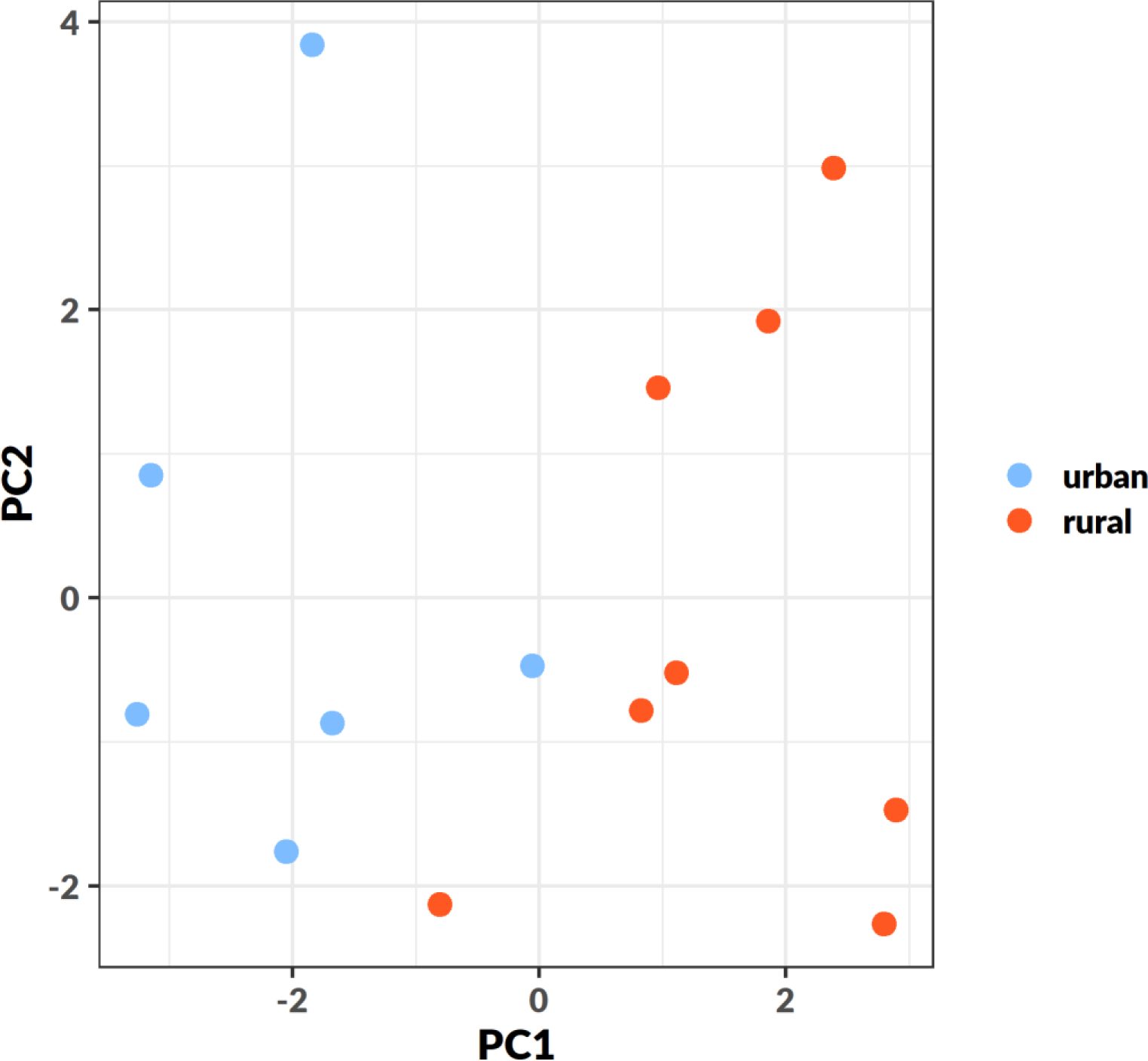
PCA biplot. PC1 explained 25.6%, and PC2 20.4% of the total variation in response variables. Sites were separated along PC1.

## Discussion

In general, serum parameters at both sites were within published ranges for grey squirrels (23,34) and reference ranges for another member of *Sciuridae*, Prevost’s squirrel (*Callosciurus prevostii*) (35) (Figs. 1, 2, 3). We found no difference in body condition between sites, therefore the chemistry values in the present study were not directly related to body condition as measured here (size-corrected weight). Even if serum analytes are not correlated with body condition, we can begin to understand some potential differences in habitat quality between our urban and rural sites, and how this affects the nutritional status of these populations with these data.

We found elevated blood glucose in urban squirrels, supporting our expectation that individuals with easy access to human food waste have higher sugar intake. This result was consistent with previous findings in raccoons and baboons (18, 20), potentially reflecting a more general trend among urban mammalian omnivores. In addition to increased blood sugar, chloride, phosphorus, and potassium concentrations varied in ways that are consistent with metabolic disorders, which in humans frequently co-occur with electrolyte abnormalities (36). However, these results might reflect differing compositions of anthropogenic versus natural foods, rather than physiological responses to a change in diet (37). Given similar body conditions between urban and rural sites, and that glucose values were within previously published ranges, it is currently unclear if these squirrels are at risk of developing metabolic disorders. Other results were inconsistent with known symptoms of diabetes and metabolic disorders in humans. We found lower ALT in urban populations, and no differences in other liver enzymes – human diabetic and pre-diabetic states are characterized by high levels of ALT, AST, and GGT (38, 39). Due to their ecology and physiology, grey squirrels may have mechanisms in place for regulating body mass and fat accumulation which reduce the risk of metabolic disturbance while accumulating lipid reserves. Grey squirrels are a seasonal fat-accumulating species that are relatively inactive during winter, but do not hibernate (23). They have high metabolic rates and calorie requirements, and are efficient users of available energy (40, 41). Peak annual mass is reached in autumn: during this time squirrels are hyperphagic, and carry more adipose tissue (42, 43). Captive grey squirrels also displayed voluntary reduction in food intake during winter periods (41). Thus, increased access to supplemental food may help autumn weight gain in grey squirrels—similar to brown bears, which are better able to optimize food intake to maximize weight gain prior to hibernation when they have access to anthropogenic food (14, 44). These characteristics suggest that grey squirrels may be pre-adapted to exploit transient food excesses, such as human food wastes, and therefore are robust to major health consequences of urban diets.

We also note interesting patterns with regard to markers for serum protein. Serum albumin reflects long-term protein status, while urea levels shift in response to short-term protein availability (45). Creatinine is reduced when muscle mass decreases due to protein catabolism in the body (45, 46). Although we did not find statistically detectable differences between populations in creatinine and urea, in both cases they tended to be lower in urban populations. Total protein was not different between sites and was on the high margin compared to previously published intervals (23, 34), indicating that neither population was malnourished, at least with respect to protein intake. Serum albumin was comparable in urban and rural squirrels, and though globulin trended higher in urban squirrels, the difference was small. However, we found significantly lower ratios of albumin to globulin (A:G ratios) in urban squirrels. Globulin is expected to be increased after infection, and is also positively associated with inflammatory responses and nutritional stress (47). Individuals in better physiological condition have higher A:G ratios. This result is interesting but difficult to interpret at this stage, because inflammation is typically the first physiological response to a stressor of any kind. Urbanization introduces numerous stressors which might cause inflammation, such as pollution, increased disease transmission, and dietary changes (48, 49). Obesity and metabolic disorders are also characterized by a chronic pro-inflammatory state (50). Previous studies using transcriptomic data report candidate genes associated with metabolism and immunity which are under selection in cities, and greater expression of genes involved in the inflammatory response in urban populations (51, 52). Our results align with general trends of greater physiological stress associated with environmental stressors in urban populations, but the underlying causes remain to be explored.

There are few published studies of serum biochemistry in grey squirrels, making inferences about individual health difficult. Our values were well within reported ranges in Hoff et al. (34) (Figs. 1, 2, 3). We note however, that those ranges were obtained from samples taken year-round, and thus might obscure potential seasonal variation in analyte levels. Autumn hyperphagia and rapid weight gain would be expected to elevate serum glucose and lipid markers including cholesterol and albumin, which in mammals is involved in fatty acid transport via the circulatory system (53). Seasonal variation in diet and food intake in grey squirrels (23) would also drive seasonal differences in blood chemistry, making comparison of our values to annual averages in serum parameters not ideal.

Our main findings—higher glucose, shifts in electrolyte balance, and lower A:G ratio—suggest physiological responses to differing habitat quality between urban and rural sites. Taken together, this is evidence that urban grey squirrels are less healthy than rural ones in our study populations. We note that this was an exploratory analysis, and results should be interpreted with caution. Sample sizes were small, and many of the variables examined here can vary based on a number of factors such as age, individual health, time since last meal, and stress associated with capture and handling. Here, all squirrels were baited, captured and handled in the same way, in essence controlling for these effects on serum biochemistry. However, in future the effects of stress and food intake on glucose might be minimized by measuring glycated serum proteins (as in (20)), which is an indicator of circulating blood sugar integrated over a longer period of time rather than instantaneous measurements of serum glucose concentrations. Of note, interpretation of glycated serum proteins on needs to be done with caution because values will be a function of integrated circulating carbohydrate levels and the relative turnover rates of circulating proteins. Albumin is a predominant soluble extracellular protein in mammal blood and based on the half-life of albumin (~ 17-20 days for humans, ~ 1.5-3 days for rats (54–57)) it is likely there is a substantial interspecific size-scaling effect where the period of integration of circulating carbohydrate levels based on glycated serum proteins will be much shorter for smaller species. Future studies examining the risk of metabolic disorders in grey squirrels could additionally measure circulating hormone levels, such as insulin, and urine glucose to encompass the main symptoms used to diagnose metabolic disorders in other species (11, 12, 58). Furthermore, pinpointing causes of inflammation would be informative for identifying specific features of urbanization that have physiological effects on wildlife. Potential sources of inflammation could be untangled using hematological tests to distinguish between infection (parasitic, viral, or bacterial), or other chronic influences (e.g., diet, pollution). We note that we obtained blood smears for urban individuals and a small subset of the rural individuals presented here (*n* = 2), but due to lack of rural samples were unable to make a comparison. However, among those individuals for which we did obtain samples, no blood parasites were detected.

The effects of food provisioning on individual health has important impacts on population and evolutionary dynamics. Food subsidies are partly responsible for increased abundances and high population densities seen in successful urban species like rats, white-tailed deer, raccoons, and grey squirrels (25, 59). However, as our results demonstrate, greater abundance does not always correspond to good overall health (20, 60, 61), meaning large urban populations may suffer higher mortality. Food provisioning can have indirect consequences on survival by promoting the spread of disease and parasites in denser populations (3, 62) which are weakened by poor nutrition and stress. In this way, supplemental food may be an important selection pressure in cities shaping evolution in successful urban species. Higher prey density in cities, in addition to supplemental human food, also increases the frequency of human-wildlife interactions (44, 60, 61, 63). What this means for population demographics in the long-term—even for species which do well in cities—is unclear. The consequences of human food waste reverberate throughout levels of the urban ecosystem, and understanding these complex relationships is important for both safety and controlling populations of synanthropic species in cities.

## Acknowledgements

CS, RPK, and CJG were supported by a Natural Sciences and Engineering Research Council of Canada Discovery Grant to CJG. CS and RPK were additionally supported by U. Manitoba Graduate Fellowships and a U. Manitoba Graduate Enhancement of Tri-council funding grant to CJG. Contribution by JRT was supported by the Canada Research Chairs program. We would like to thank Leah Kathan and Alyssa Garrard for their fieldwork contributions. We are grateful to Neil Pople, Rhonda Gregoire, Amanda Salo, and the Manitoba Veterinary Diagnostics Services lab for their assistance with collecting and processing serum samples.

## References

1. Murray M, et al. (2019) City sicker? A meta-analysis of wildlife health and urbanization. Front Ecol Environ In Press:1–9.

2. Birnie-Gauvin K, Peiman KS, Gallagher AJ, De Bruijn R, Cooke SJ (2016) Sublethal consequences of urban life for wild vertebrates. Environ Rev 24(4):416–425.

3. Oro D, Genovart M, Tavecchia G, Fowler MS, Martínez-Abraín A (2013) Ecological and evolutionary implications of food subsidies from humans. Ecol Lett 16(12):1501–1514.

4. Plummer KE, Bearhop S, Leech DI, Chamberlain DE, Blount JD (2013) Winter food provisioning reduces future breeding performance in a wild bird. Sci Rep 3. doi:10.1038/srep02002.

5. Weaver CM, et al. (2014) Processed foods: contributions to nutrition. Am J Clin Nutr 19(1525-42):61–68.

6. McKinney ML (2002) Urbanization, Biodiversity, and Conservation. Bioscience 52(10):883–890.

7. Mendoza JA, Drewnowski A, Christakis DA (2007) Dietary energy density is associated with obesity and the metabolic syndrome in U.S. adults. Diabetes Care 30(4):974–979.

8. Eckel RH, Alberti KGMM, Grundy SM, Zimmet PZ (2010) The metabolic syndrome. Lancet 375(9710):181–183.

9. Heidemann C, et al. (2008) Dietary patterns and risk of mortality from cardiovascular disease, cancer, and all causes in a prospective cohort of women. Circulation 118(3):230–237.

10. Lund EM, Armstrong PJ, Kirk CA, Klausner JS (2006) Prevalence and risk factors for obesity in adult dogs from private US veterinary practices. Int J Appl Res Vet Med 4(2):177–186.

11. Greco DS (2001) Diagnosis of diabetes mellitus in cats and dogs. Vet Clin North Am Small Anim Pract 31(5):845–853.

12. Ciobotaru E (2013) Spontaneous Diabetes Mellitus in Animals. Diabetes Mellitus - Insights and Perspectives (InTech), pp 271–296.

13. Townsend AK, Staab HA, Barker CM (2019) Urbanization and elevated cholesterol in American Crows. Condor 121(3):1–10.

14. Coogan SCP, Raubenheimer D, Stenhouse GB, Coops NC, Nielsen SE (2018) Functional macronutritional generalism in a large omnivore, the brown bear. Ecol Evol 8(4):2365–2376.

15. Klimentidis YC, et al. (2011) Canaries in the coal mine: a cross-species analysis of the plurality of obesity epidemics. Proc R Soc Biol Sci 278(1712):1626–32.

16. Cypher BL, Frost N (1999) Condition of San Joaquin Kit Foxes in Urban and Exurban Habitats. J Wildl Manage 63(3):930–938.

17. Harrison RL (1997) A Comparison of Gray Fox Ecology between Residential and Undeveloped Rural Landscapes. J Wildl Manage 61(1):112–122.

18. Banks WA, Altmann J, Sapolsky RM, Phillips-Conroy JE, Morley JE (2003) Serum leptin levels as a marker for a syndrome X-like condition in wild baboons. J Clin Endocrinol Metab 88(3):1234–1240.

19. Harveson PM, Lopez RR, Collier BA, Silvy NJ (2007) Impacts of urbanization on Florida Key deer behavior and population dynamics. Biol Conserv 134(3):321–331.

20. Schulte-Hostedde AI, Mazal Z, Jardine CM, Gagnon J (2018) Enhanced access to anthropogenic food waste is related to hyperglycemia in raccoons (Procyon lotor). Conserv Physiol 6(1):1–6.

21. Hanks J (1981) Characterization of Population Condition. Dynamics of Large Mammal Populations, eds Fowler CW, Smith TD (John Wiley & Sons), pp 47–73.

22. Bertolino S (2009) Animal trade and non-indigenous species introduction: The world-wide spread of squirrels. Divers Distrib 15(4):701–708.

23. Koprowski JL (1994) Sciurus carolinensis. Mamm Species (480):1–9.

24. Bonnington C, Gaston KJ, Evans KL (2014) Squirrels in suburbia: Influence of urbanisation on the occurrence and distribution of a common exotic mammal. Urban Ecosyst 17(2):533–546.

25. Parker TS, Nilon CH (2008) Gray squirrel density, habitat suitability, and behavior in urban parks. Urban Ecosyst 11(3):243–255.

26. Mahdawi A (2018) The new Pizza Rat? New York squirrel filmed snacking on avocado. Guard. Available at: https://www.theguardian.com/us-news/2018/jun/08/avocado-squirrel-pizza-rat-new-york.

27. Vinopal C (2017) Look Out, Pizza Rat…Say Hello to Pizza Squirrel. Washingtonian. Available at: https://www.washingtonian.com/2017/08/22/look-pizza-rat-say-hello-pizza-squirrel/.

28. r/FatSquirrels (2019) Available at: https://reddit.com/r/fatsquirrels.

29. r/SquirrelsEatingPizza (2019) Available at: https://reddit.com/r/squirrelseatingpizza.

30. Koprowski JL (2002) Handling tree squirrels with a safe and efficient restraint. Wildl Soc Bull 30(1):101–103.

31. Schulte-Hostedde AI, Zinner B, Millar JS, Hickling GJ (2005) Restitution of mass-size residuals: Validating body condition indices. Ecology 86(1):155–163.

32. Schulte-Hostedde AI, Millar JS, Hickling GJ (2001) Evaluating body condition in small mammals. Can J Zool 79(6):1021–1029.

33. R Core Team (2013) R: A Language and Environment for Statistical Computing. Available at: http://www.r-project.org/.

34. Hoff GL, McEldowny LE, Bigler WJ, Kuhns LJ, Tomas JA (1976) Blood and urinary values in the Gray Squirrel. J Wildl Dis 12:349–352.

35. J A Teare ed. (2013) Callosciurus_prevostii_No_selection_by_gender__All_ages_combined_Standard_International_Units 2013_CD.html in ISIS Physiological Reference Intervals for Captive Wildlife: A CD-ROM Resource.

36. Liamis G, Liberopoulos E, Barkas F, Elisaf M (2014) Diabetes mellitus and electrolyte disorders. World J Clin Cases 2(10):488.

37. Choi CS, Lee FN, McDonough AA, Youn JH (2002) Independent regulation of in vivo insulin action on glucose versus K + uptake by dietary fat and K + content. Diabetes 51(4):915–920.

38. Music M, et al. (2015) Metabolic Syndrome and Serum Liver Enzymes Level at Patients with Type 2 Diabetes Mellitus. Med Arch (Sarajevo, Bosnia Herzegovina) 69(4):251–255.

39. Ahn HR, et al. (2014) The association between liver enzymes and risk of type 2 diabetes: The Namwon study. Diabetol Metab Syndr 6(1):2–9.

40. Montgomery SD, Whelan JB, Mosby HS (1975) Bioenergetics of a Woodlot Gray Squirrel Population. J Wildl Manage 39(4):709–717.

41. Ludwick RL, Fontenot J, Mosby HS (1969) Energy Metabolism of the Eastern Gray Squirrel. J Wildl Manage 33(3):569–575.

42. Knee C (1983) Squirrel energetics. Mamm Rev 13(2-4):113–122.

43. Pereira ME, Aines J, Scheckter JL (2002) Tactics of Heterothermy in Eastern Gray Squirrels (Sciurus Carolinensis). J Mammal 83(2):467–477.

44. Coogan SCP, Raubenheimer D (2016) Might macronutrient requirements influence grizzly bear-human conflict? Insights from nutritional geometry. Ecosphere 7(1):1–15.

45. Caldeira RM, Belo AT, Santos CC, Vazques MI, Portugal A V. (2007) The effect of long-term feed restriction and over-nutrition on body condition score, blood metabolites and hormonal profiles in ewes. Small Rumin Res 68(3):242–255.

46. Robert KA, Schwanz LE (2013) Monitoring the health status of free-ranging tammar wallabies using hematology, serum biochemistry, and parasite loads. J Wildl Manage 77(6):1232–1243.

47. Kilgas P, Mänd R, Mägi M, Tilgar V (2006) Hematological parameters in brood-rearing great tits in relation to habitat, multiple breeding and sex. Comp Biochem Physiol - A Mol Integr Physiol 144(2):224–231.

48. Isaksson C (2015) Urbanization, oxidative stress and inflammation: A question of evolving, acclimatizing or coping with urban environmental stress. Funct Ecol 29(7):913–923.

49. Bradley CA, Altizer S (2007) Urbanization and the ecology of wildlife diseases. Trends Ecol Evol 22(2):95–102.

50. Hotamisligil GS (2006) Inflammation and metabolic disorders. Nature 444(7121):860–867.

51. Harris SE, Munshi-South J, Obergfell C, Neill RO (2013) Signatures of Rapid Evolution in Urban and Rural Transcriptomes of White-Footed Mice (Peromyscus leucopus) in the New York Metropolitan Area. PLoS One 8(8):1–19.

52. Watson H, Videvall E, Andersson MN, Isaksson C (2017) Transcriptome analysis of a wild bird reveals physiological responses to the urban environment. Sci Rep 7:1–10.

53. LeBlanc PJ, et al. (2001) Correlations of plasma lipid metabolites with hibernation and lactation in wild black bears Ursus americanus. J Comp Physiol - B Biochem Syst Environ Physiol 171(4):327–334.

54. Urlich F, Tarver H, Li CH (1954) Effects of growth and adrenocorticotropic hormones on the metabolism of albumin in hypophysectomized rats. J Biol Chem 209(1):117–25.

55. Allison AC (1960) Turnovers of erythrocytes and plasma proteins in mammals. Nature 188(4744):37–40.

56. Day JF, Thornburg RW, Thorpe SR, Baynes JW (1979) Nonenzymatic glucosylation of rat albumin: Studies in vitro and in vivo. J Biol Chem 254(19):9394–9400.

57. Schultze HE, Heremans JF (1966) Molecular Biology of Human Proteins, with Special Reference to Plasma Proteins. Volume 1: Nature and Metabolism of Extracellular Proteins. (Elsevier, Amsterdam).

58. Heidt GA, Conaway HH, Frith C, Farris Jr. HE (1984) Spontaneous Diabetes mellitus in a captive Golden-manted ground squirrel, Spermophilus lateralis (Say). J Wildl Dis 20(3):253–255.

59. Adams LW (1994) Urban wildlife habitats: A landscape perspective (University of Minnesota Press).

60. Fedriani JM, Fuller TK, Sauvajot RM (2001) Does availability of anthropogenic food enhance densities of omnivorous mammals? An example with coyotes in southern California. Ecography (Cop) 24(3):325–331.

61. Murray M, et al. (2015) Greater consumption of protein-poor anthropogenic food by urban relative to rural coyotes increases diet breadth and potential for human-wildlife conflict. Ecography (Cop) 38(12):1235–1242.

62. Orams MB (2002) Feeding wildlife as a tourism attraction: a review of issues and impacts. Tour Manag 23:281–293.

63. Thiemann GW, Stahl RS, Baruch-Mordo S, Breck SW (2008) Trans fatty acids provide evidence of anthropogenic feeding by black bears. Human-Wildlife Interact 2(2):183–193.

